# Every Piece of the Puzzle Matters: A Novel Zinc-Binding Site in the Luminal Domain of STIM1 Drives Clustering

**DOI:** 10.1101/2025.06.12.659323

**Authors:** Benedicte Alary, Viktoriia E. Baksheeva, Sabrina Beaumier, Géraldine Ferracci, Claude Villard, Andrey V. Golovin, Sébastien Courrier, François Devred, Marc Bartoli, Philipp O. Tsvetkov

## Abstract

STIM1 is pivotal in the tightly regulated mechanism controlling calcium homeostasis in the ER. It is activated by calcium dissociation from its EF-hand domain when ER calcium levels decrease, leading to its interaction with the ORAI channel to initiate calcium influx. Despite advancements in understanding the complex STIM1 machinery, including its domain organization and structural rearrangements upon activation, many aspects of this process remain poorly understood. In this study, we focused on a small conserved region situated upstream to the EF-hand, which has been previously shown to be involved in the modulation of STIM1 by ROS. Our findings reveal that this region binds zinc and plays a pivotal role in STIM1 activation by promoting its clustering, a process essential for the activation of calcium influx. These results revealed the functional importance of this domain and added a crucial piece to the puzzle of how calcium and zinc signaling are interconnected.

**Graphical abstract:** 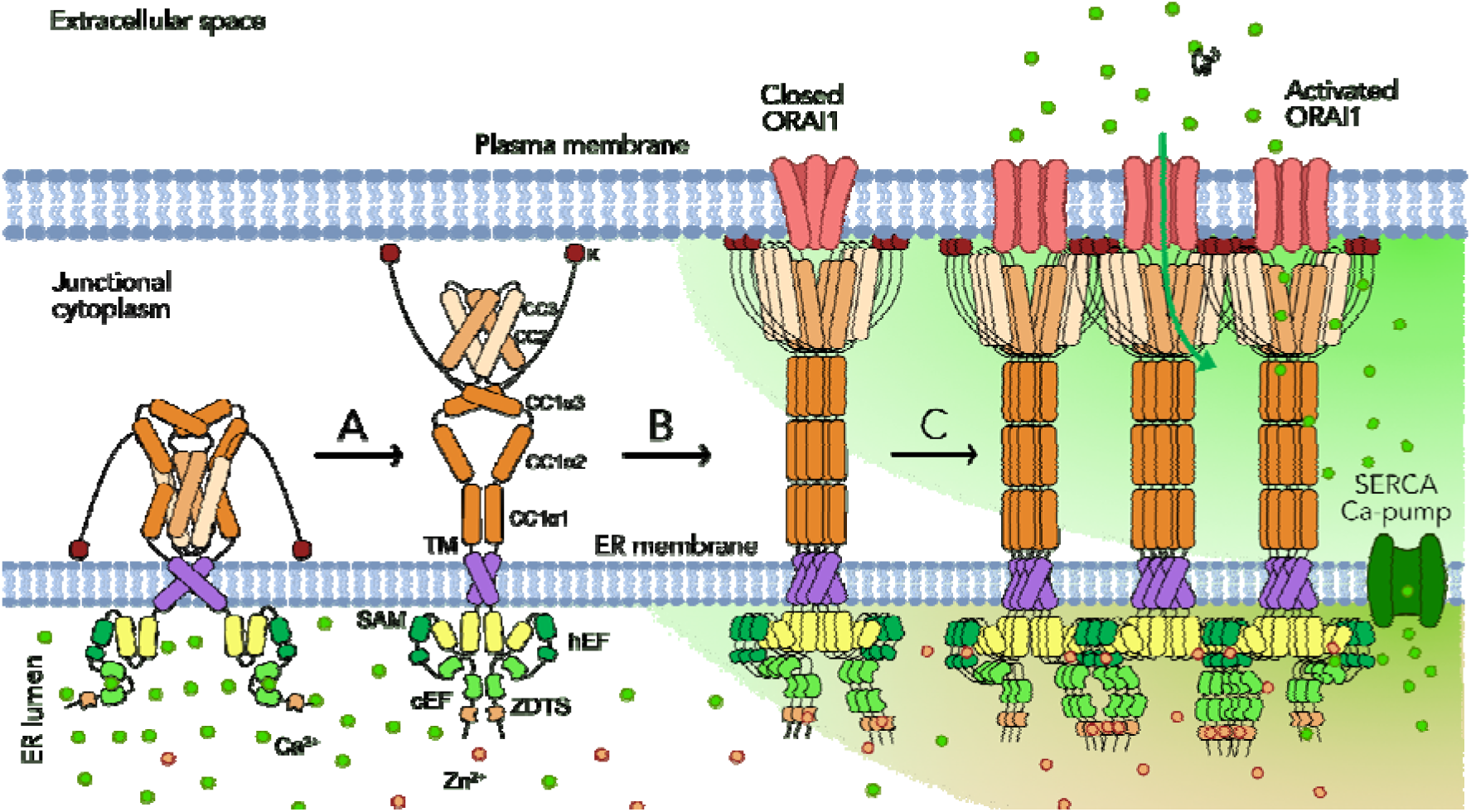

## Introduction

Calcium signaling is essential for a wide array of cellular functions, including secretion, excitation, contraction, metabolism, transcription, growth, and apoptosis^1^. The endoplasmic reticulum (ER) plays a central role in these processes as an intracellular calcium store, enabling rapid calcium signaling and providing an optimal environment for protein folding and trafficking. However, disruptions in calcium homeostasis, particularly a reduction in luminal calcium levels in the ER, can lead to cellular stress and contribute to various pathologies ^2^.

To restore calcium balance, cells rely on store-operated calcium entry (SOCE), a tightly regulated mechanism mediated by two key protein families: stromal interaction molecule (STIM) and calcium release-activated calcium channel protein (ORAI). These proteins form calcium release-activated channels (CRACs), where STIM proteins detect ER calcium depletion and activate ORAI calcium channels on the plasma membrane. This interaction triggers a calcium influx into the cytosol. Then the sarco/endoplasmic reticulum calcium ATPase (SERCA) pump actively transports calcium back into the ER to replenish its stores, ensuring cellular calcium equilibrium.

STIM proteins, predominantly STIM1 and STIM2, are single-pass transmembrane proteins primarily localized to the ER membrane, though they are also found in lysosome-related organelles and the plasma membrane ^3,4^ . Both homologs share a conserved luminal domain responsible for calcium sensing, while their cytosolic domains, which interact with ORAI channels, display greater variability ^5–8^. Alternative splicing generates isoforms such as STIM1L, which is muscle-specific, and STIM1A, an inhibitor of SOCE. The most widely studied isoform, STIM1, consists of 685 amino acids and is ubiquitously expressed.

STIM1 and the SOCE mechanism are critical for physiological processes such as immune cell activation and muscle function. Mutations in the STIM1 gene are associated with diverse pathologies. Loss-of-function mutations lead to CRAC channelopathy, characterized by muscular hypotonia, ectodermal dysplasia, severe combined immunodeficiency (SCID)-like conditions, and autoimmunity, while gain-of-function mutations are linked to tubular aggregate myopathy (TAM)^9^ and Stormorken syndrome, both of which manifest as muscle weakness ^10^. STIM1 and SOCE have also been shown to play an important role in cancer progression. Increased expression of the STIM and ORAI proteins and SOCE activity is generally associated with elevated proliferation, migration and invasion of cancer cells^11^.

Similar to all other isoforms in the family, STIM1’s intraluminal region contains two EF-hand domains and a SAM (sterile alpha motif) domain (Fig.1A,B). It has been proposed that the canonical EF-hand domain (cEF) actively binds calcium, while the non-canonical EF-hand or hidden EF-hand (hEF) stabilizes this binding. The exact number of calcium ions bound to STIM1 remains debated ^6,7,12,13^. Overall, the EF-hands bind and detect calcium levels, and the SAM domain promotes oligomerization of the protein upon activation.

**Figure 1.**
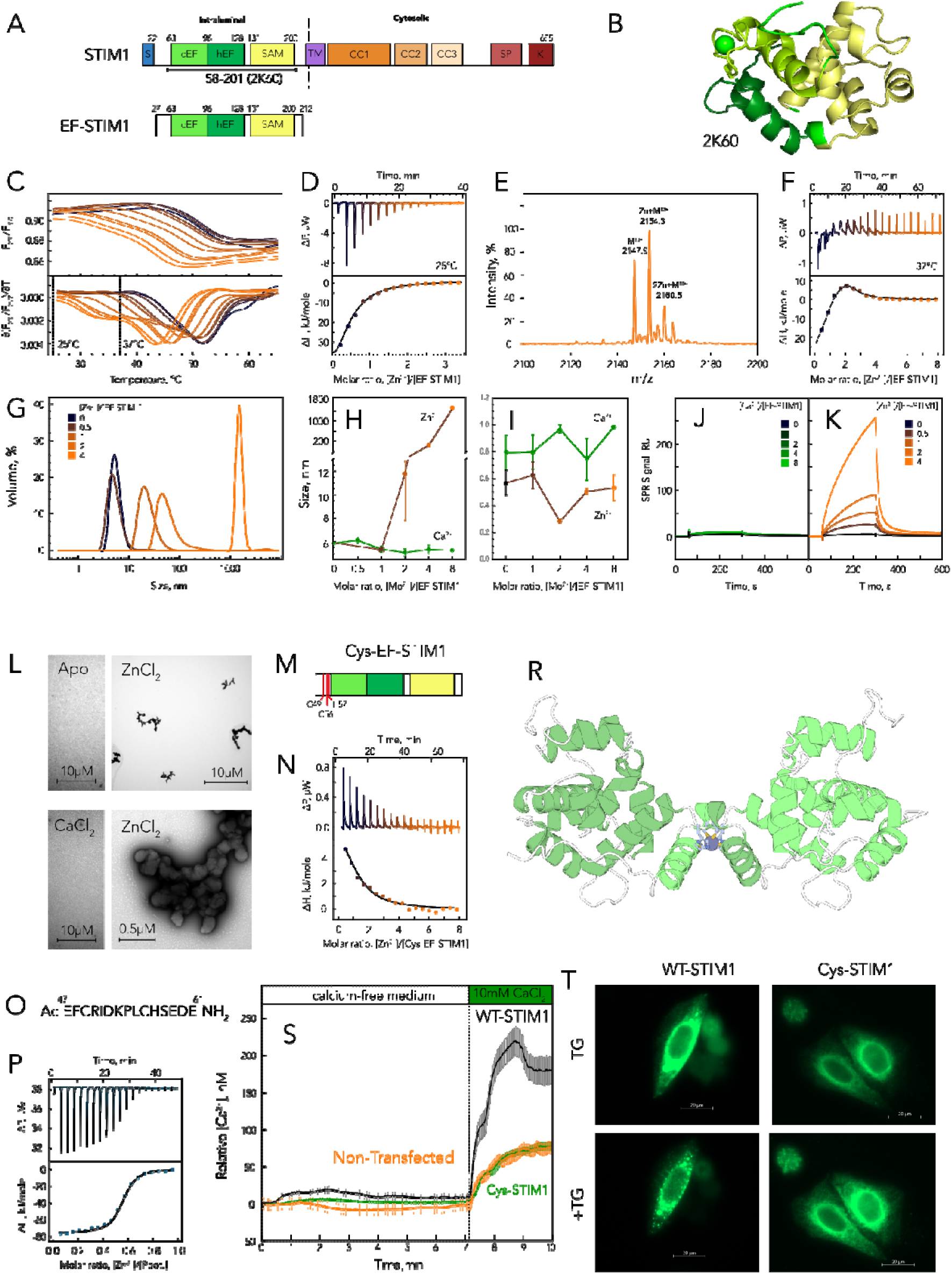
In vitro analysis of zinc binding to the luminal domain of STIM1 and the effects of zinc-binding disruptive mutations on STIM1 activation in cells. (A) Domain organization of STIM1 and EF-STIM1 a 27-212 a.a. fragment used in the study. (B) Structure of the luminal domain of STIM1 in complex with one calcium ion (PDBID: 2K60)^7^. (C) Denaturation curves of EF-STIM1 in the presence of an increasing concentration of zinc obtained by nanoDSF and their derivatives. (D,F) Titration curve representing EF-STIM1 binding with zinc ions and its fitting at 25°C and 37°C respectively. (E) Mass-spectra of Zn^2+^-bound EF-STIM1. (G) Size distribution by volume obtained by DLS for EF-STIM1 in the presence of increasing concentration of zinc ions. (H) Size of most abundant species in the presence of calcium and zinc ions. (I) Polydispersity index of EF-STIM1 in the presence of calcium and zinc ions. (J, K) SPR assay of EF-STIM1 self-association in the presence of increasing concentrations of calcium and zinc respectively. (L) TEM images of zinc-induced aggregates. (M) Mutations in the zinc-binding site of EF-STIM1. (N) ITC titration curve representing Cys-STIM1 binding with zinc ions and its fitting at 37°C. (O) Primary sequence of peptide corresponding to high-affinity zinc-binding site. (P) ITC titration curve representing zinc ions to peptide and its fitting with one-set-of-sites model. (R) Structure of EF-STIM1 obtained using ML approach. (S) Relative calcium concentration in HEK cells non-transfected or transfected with WT-STIM1 or Cys-STIM1 fused to GFP. Cells loaded with fura-2-red AM in a calcium-free medium were treated with TG and CaCl_2_ was added to the medium. (T) Fluorescence microscopy on HeLa cells transfected with WT-STIM1 or Cys-STIM1 fused to GFP before (top panel) and 10 min after the addition of TG (lower panel) at 37°C

The activation of STIM1 and SOCE is a complex, multi-stage process^14^. Despite substantial progress in understanding the underlying mechanisms, a comprehensive and detailed description of this process remains elusive. Current models propose that, at rest, when ER calcium stores are saturated, the EF-hand domains of STIM1 remain bound to calcium ions. In this state, the cytosolic regions of two STIM1 monomers are closely aligned, adopting a compact, self-folded conformation, while their transmembrane domains are tightly paired, forming a stable resting dimer. In contrast, the luminal domains of the two STIM1 subunits remain monomeric, with the EF-hand and SAM domains folding cooperatively into a stable conformation. When calcium stores are depleted, calcium release from the cEF-hand triggers the unfolding of the EF-SAM domain including a significant drop in alpha-helical content. This unfolding exposes hydrophobic surfaces that probably promote dimerization and/or oligomerization of the luminal portion of the protein ^6,7^, leading to a conformational change that propagates to the transmembrane domain and cytosolic regions ^14^. In the absence of intraluminal calcium, the activated STIM1 oligomerizes and relocates to junctional zones between the ER/SR and the plasma membrane to interact with and activate the ORAI1 calcium channels ^15^.

Preceding the cEF-hand domain, there is a 60-amino-acid-long region with an unresolved structure and poorly understood functions. This region includes a small, highly conserved sequence across different species, which contains two closely positioned and even more highly conserved cysteine residues, Cys49 and Cys56. Here, we provide strong evidence that Cys49 and Cys56 could also be implicated in the zinc-dependent regulation of STIM1 activity by favoring oligomerization of its luminal domain, which might be responsible for STIM1 clusterization in cells.

## Results

### The Intraluminal Domain of STIM1 Binds Two Zinc Ions

Zinc binding is known to profoundly influence protein structure and thermostability. To elucidate the interaction between zinc and the intraluminal domain of STIM1 (EF-STIM1), we performed temperature-induced denaturation experiments using nanoDSF, progressively increasing the concentration of zinc ions. Our findings demonstrate a gradual destabilization of the EF-STIM1 structure upon increase of zinc concentration (Fig. 1C). While zinc can bind to some proteins in a nonspecific manner, resulting in continuous destabilization, the saturation observed in the destabilization of EF-STIM1 suggests that zinc binds to distinct, specific sites on the protein. To further characterize the thermodynamic parameters of zinc binding to EF-STIM1, we employed isothermal titration calorimetry (ITC), a gold-standard method for such analyses. Titration of EF-STIM1 with zinc at 25°C revealed a single, enthalpy-favorable binding site with a dissociation constant (K_d_) of 28 µM (Fig. 1D). Mass spectrometry, used to confirm stoichiometry, suggested the potential presence of two binding sites (Fig. 1E). Given that the ITC signal is entirely dependent on the enthalpy of binding, which can vary with temperature, we performed additional ITC experiments at 37°C to investigate the possibility of a second zinc-binding site. Indeed, the titration curve at this temperature exhibited biphasic behavior, indicative of multiple binding sites. A two-set-of-site binding model with the stoichiometry of both sites fixed to one provided an excellent fit to the data, confirming the presence of two zinc-binding sites on EF-STIM1 with K_d_ of 16 µM for entropy-driven site and 2.7 µM for enthalpy favorable site. Very close in absolute value, but opposite in sign the enthalpy of sites (36.4 and -31.0 kJ/mole), explains the relatively low resulting ITC signal. The unfavorable enthalpy of the second zinc binding as well as wide titration peaks indicative of slow processes lead us to the hypothesis that this binding is coupled with the changes in the oligomeric state of the protein.

### Zinc Modulates the Oligomeric State of the STIM1 Intraluminal Domain

To investigate the oligomeric state of the EF-STIM1 fragment, we analyzed its behavior in solution under increasing concentrations of zinc and calcium ions using dynamic light scattering (DLS) (Fig. 1G–I). In agreement with previous reports, elevating the calcium concentration led to a modest reduction in the hydrodynamic radius of the predominant EF-STIM1 species—from approximately 6□nm to about 5□nm at a twofold molar excess of Ca²□ (Fig. 1H). This size reduction approximates that of a known smaller Ca²□-bound fragment of the intraluminal domain (□4□nm; Fig. 1B), with the slightly larger dimensions observed here being consistent with the use of a correspondingly larger fragment in our experiments.

Because DLS is sensitive to changes in the hydrodynamic volume of proteins, the observed decrease could stem from either dimer dissociation or a conformational transition from a partially unfolded apo-state to a more compact Ca²□-bound form. To discern between these possibilities, we employed surface plasmon resonance (SPR), which directly senses the variation of mass on the SPR chip. Notably, SPR data revealed that the presence or absence of calcium did not lead to auto-association of EF-STIM1 thus did not alter the oligomeric state of EF-STIM1. This finding strongly suggests that the reduction in hydrodynamic radius—from 6□nm to 5□nm—is attributable to protein folding rather than to the dissociation of dimers.

In contrast to calcium, zinc exerted a dramatic influence on the oligomeric state of the STIM1 luminal domain. DLS experiments showed that at sub-equimolar concentrations, zinc initially induced a slight decrease in EF-STIM1 size, likely reflecting structural stabilization analogous to that observed with calcium. However, upon reaching equimolar zinc concentrations, EF-STIM1 assembled into 20□nm oligomers. With a twofold molar excess of zinc, even larger oligomers (40–50□nm) were detected, and saturating conditions—achieved at a fourfold zinc excess—led to the formation of aggregates ranging from 1 to 2□µm in diameter. Interestingly, although the polydispersity index briefly increased at a 0.5-fold zinc excess, it subsequently decreased and reached a minimum at a twofold excess, indicating that the oligomers formed under these conditions were homogeneous in size. Consistent with the DLS results, SPR measurements demonstrated a progressive increase in STIM1 auto-association as the zinc concentration was raised.

To visualize the structure of EF-STIM1 oligomers, we employed transmission electron microscopy (TEM) (Fig. 1L). In the absence of metal ions, as well as in the presence of calcium, no visible structures were observed. However, with a fourfold excess of zinc, we detected aggregates whose size closely matched our DLS data. A detailed examination of these aggregates revealed an internal structure composed of rounded particles, consistent in size with the oligomers observed in the DLS measurements.

### Cysteine Zinc-Binding Site Responsible for Modulating the Oligomeric State of STIM1

EF-hand motifs are known to bind zinc ions, often inducing conformational changes similar to those observed with calcium. In the case of EF-STIM1, this is likely true as well, given the slight compaction of the protein structure observed at low concentrations of both calcium and zinc. Therefore, we hypothesize that the binding of a second zinc ion could be responsible for alterations in the oligomeric state of EF-STIM1. Since zinc coordination is most commonly mediated by cysteine and histidine residues, and these residues are clustered immediately before the first EF-hand domain, we propose that they form a high-affinity zinc-binding site. To test this hypothesis, we generated an EF-STIM1 mutant in which key cysteine Cys49, Cys56 and histidine His57 residues were substituted with alanine (Fig. 1M). In parallel, we synthesized a peptide corresponding to residues 47–61 (Fig. 1O), which includes all potential zinc-chelating amino acids.

Isothermal titration calorimetry (ITC) experiments provided clear support for our hypothesis. Titration of the EF-STIM1 mutant with zinc ions (Fig. 1N) revealed a single enthalpy-driven binding event with a stoichiometry of approximately 1.1 and a dissociation constant (K_d_) of 32 µM—consistent with the low-affinity zinc-binding site observed in wild-type EF-STIM1, likely located within one of the EF-hands. In contrast, ITC analysis of the peptide (Fig. 1P) showed a high-affinity binding event with a K_d_ of 0.6 µM. Notably, the binding stoichiometry for the peptide was around 0.5, suggesting the formation of zinc-linked peptide dimers. These results highlight the functional role of the high-affinity zinc-binding site, mediated by cysteine residues, in modulating the oligomeric state of STIM1. Interestingly, an *in silico* model of the zinc-linked EF-STIM1 dimer, obtained using a machine learning (ML) approach (Fig. 1J), allowed us to hypothesize that this dimerization does not occur between the luminal domains of an STIM1 dimer but rather links two STIM1 dimers, thus promoting cluster formation.

To explore the functional significance of this site in cells, we transfected HeLa and HEK cells with either full-length wild-type STIM1 or a mutant version with modified cysteine and histidine residues (Cys-STIM1). We then assessed its impact on SOCE activation by inducing conditions that trigger SOCE using thapsigargin (TG), a well-established inhibitor of the ER Ca^2+^ ATPase SERCA. TG depletes ER Ca^2+^ stores, activating STIM1. The addition of TG to a calcium-free medium induces the release of ER Ca^2+^ into the cytoplasm, initiating STIM1 activation. Subsequent injection of calcium into the extracellular medium triggers calcium entry via ORAI1 CRAC channels (Fig. 1S). In untransfected HEK cells, endogenous STIM1 proteins are present and functional, leading to an approximate increase in calcium concentration of 76 nM. In cells transfected with WT-STIM1 constructs, the overexpression of active STIM1 significantly enhances SOCE activation, leading to cytosolic calcium concentrations rising to as high as 220 nM after calcium addition. In contrast, transfection with Cys-STIM1 constructs produces calcium entry levels comparable to those observed in the untransfected condition, with a calcium peak of approximately 78 nM after SOCE activation. Moreover, the area under the curve after calcium addition further highlights these differences: 168.6±53.1 mmol·L^-1^·min^-1^ for the untransfected condition, 575.2±74.0 mmol·L^-1^·min^-1^ for WT-STIM1, and 192.8±23.1 mmol·L^-1^·min^-1^ for Cys-STIM1. These findings indicate that overexpression of the Cys-STIM1 mutant does not enhance SOCE activity, unlike WT-STIM1, demonstrating that Cys-STIM1 represents an inactive form of the protein.

Our *in vitro* experiments demonstrate that zinc binding to this site induces oligomerization of the intraluminal domain of STIM1. Therefore, we hypothesize that the lack of additional calcium influx observed in Cys-STIM1 transfected cells could be attributed to the inability of Cys-STIM1 to undergo oligomerization. We transfected HeLa cells with WT and Cys-mutated STIM1 coupled with GFP, to check this hypothesis. As expected, SOCE activation by TG leads to the formation of STIM1 clusters visible in the WT-transfected cells (Fig.1T, top panels). In contrast, in the Cys-STIM1 transfected cells, TG does not lead to a redistribution of STIM1, and the proteins do not appear to form clusters (Fig.1T, bottom panels).

## Discussion

STIM1 is a pivotal calcium sensor in the endoplasmic reticulum (ER), playing a central role in store-operated calcium entry (SOCE) by activating ORAI1 upon ER calcium depletion^16^. Its complex multi-domain architecture has been the focus of extensive research over the past two decades, with numerous studies aiming to elucidate the molecular mechanisms governing its activation and structural dynamics. While the exact sequence of events in STIM1 activation remains unresolved, a recent review by Isabella Derler team has summarized the currently available key pieces that contribute to solving the puzzle of STIM1 activation^17^. Our findings add a crucial piece to this puzzle by shedding light on a small, conserved region of the luminal domain that has largely escaped the attention of most researchers, allowing us to identify another key player in the STIM1 activation cascade: the zinc ion.

Zinc’s impact on SOCE has been reported previously, though its effects were not directly attributed to STIM1. Notably, zinc ions have been shown to interact with the Orai1 channel^18^, leading to SOCE inhibition and modulation of intracellular Ca²□ oscillations^19^. We hypothesized that, similar to many Ca²□-binding proteins, STIM1 could be directly regulated by zinc. To investigate this, we focused on a small conserved region in the luminal domain of STIM1 containing two cysteine residues (Cys49 and Cys56) and one histidine—amino acids commonly involved in zinc coordination. Previous studies demonstrated that STIM1 activation can be modulated via Cys49 and Cys56, either via S-glutathionylation of Cys56 in response to oxidative stress^20^ or through disulfide bond rearrangement facilitated by the ER-resident oxidoreductase ERp57^21^. Notably, activity of some ER-resident oxidoreductase such as ERp44^22^ and PIDA1^23^ is regulated by zinc ions. Our findings strongly suggest that Cys49 and Cys56 cysteine residues also serve as key zinc-binding sites, promoting dimerization of the STIM1 luminal domain, its following oligomerization, and clustering—critical processes required for SOCE activation, challenging the traditional idea that STIM1 activation is exclusively dependent on calcium.

Indeed, our ITC data demonstrated that the luminal domain of STIM1 has two zinc binding sites with significantly different dissociation constants (K_d_) of 16 µM and 2.7 µM. Due to low overall ITC signal it was impossible to unambiguously determine the stoichiometry of zinc binding using this method. MS experiments confirmed the presence of two zinc binding sites on EF-STIM1. Subsequent ITC experiments with an EF-STIM1 cysteine mutant and a peptide fragment enabled us to identify the cysteine site as the high-affinity zinc-binding site. Notably, the stoichiometry of zinc binding to the peptide corresponding to this site suggests that a single zinc ion bridges two molecules, as previously observed for other peptides^24^ and proteins^25^. Further investigation of dimerization using ML modeling revealed that the peptides can adopt multiple conformations, with cysteine and histidine residues coordinating zinc in different configurations (data not shown). If such dimerization occurs within a single STIM1 dimer, it could induce conformational changes that propagate to the cytoplasmic domain. Alternatively, if dimerization occurs between the luminal domains of two separate STIM1 dimers, it could drive sequential oligomerization, leading to the formation of extended clusters. While our ML-based modeling supports the inter-dimer configuration, which is also consistent with our cellular assays, resolving this structural mechanism definitively will probably require high-resolution NMR analysis.

Surprisingly, our SPR and DLS experiments did not detect EF-STIM1 dimerization upon calcium dissociation, as previously reported by the Stathopulos team^6^, but only a slight increase in the molecule’s size from 5 to 6 nm, which could be associated with a partial loss of its tertiary structure, as demonstrated in the same study^6^. The Ca²□-bound EF-STIM1 size we observed by DLS is in good agreement with the NMR-resolved structure of this domain^7^. However, this contradicts the previously reported <1.5 nm size of Ca²□-bound EF-STIM1^6^, which appears unrealistically small and could be attributed to artifacts in the deconvolution of correlograms—an issue we have also occasionally observed. Moreover, the minor variations in dimer concentrations observed in glutaraldehyde cross-linking experiments^6^ could also be explained by substantial conformational differences between the apo and calcium-bound forms of EF-STIM1. The observed discrepancy in the oligomeric state of apo-EF-STIM1 could also be linked to the use of extremely low, non-physiological temperatures (4–15°C) for detecting its dimerization, as was the case in the study by the Stathopulos team^6^. Considering the contradictions regarding the effect of calcium on STIM1 dimerization and its central role in the widely accepted STIM1 activation model, this question should be carefully reassessed in future studies. Here, however, our focus remains on the effect of zinc on this process.

Since we observed no significant differences in the oligomeric state of EF-STIM1 with or without calcium, we hypothesized that zinc could play a pivotal role in luminal domain dimerization. It is worth noting that, in contrast to calcium, zinc is well known for its ability to induce protein aggregation and/or oligomerization in both physiological and pathological conditions ^26–29^. Our SPR experiments demonstrated the interaction between EF-STIM1 monomers immobilized on-chip and EF-STIM1 in solution in the presence of zinc ions. Furthermore, DLS and TEM experiments confirmed the formation of size-homogeneous oligomers in the presence of zinc ions, appearing as well-defined, rounded particles, which stick together upon further increase in zinc concentration. This observation further supports the idea that zinc binding facilitates STIM1 oligomerization, potentially driving higher-order assembly and functional clustering.

Consistent with this hypothesis, our functional assays in HeLa and HEK cells revealed that the mutant with a disrupted zinc-binding site failed to cluster in response to ER calcium depletion and was unable to enhance calcium entry. These findings further reinforce the critical role of zinc-dependent clustering in STIM1 activation and introduce an additional layer of complexity in SOCE regulation, highlighting the significance of zinc-calcium cross-talk—a fundamental interplay observed in numerous cellular processes ^30,31^.

Notably, while extracellular zinc appears to inhibit ORAI1-mediated SOCE^19^, our data strongly suggest that intracellular zinc is essential for its activation. This underscores the finely tuned regulatory balance between zinc and calcium signaling, suggesting that STIM1 activation is not solely dependent on calcium depletion but also on the dynamic availability of intracellular zinc.

Thus, we propose a model in which zinc acts as a crucial regulator of STIM1 activation (Fig.2). During ER calcium depletion, intracellular zinc levels may either remain stable or increase, directly modulating STIM1 function^32,33^. Studies have reported an inverse relationship between cytosolic calcium and ER zinc levels, further supporting the idea of a dynamic interplay between these two metal ions^30,34^. Mechanistically, calcium dissociation from STIM1’s EF-hand domain induces a significant loss of its tertiary structure, increasing its conformational flexibility. This structural change could bring the zinc-binding sites of two STIM1 molecules into closer proximity, allowing zinc to act as a bridging element between them. Zinc-induced oligomerization of STIM1 may, in turn, enhance its clustering and interaction with ORAI1, thereby promoting calcium influx into the cytosol and sustaining SOCE.

**Figure 2.**
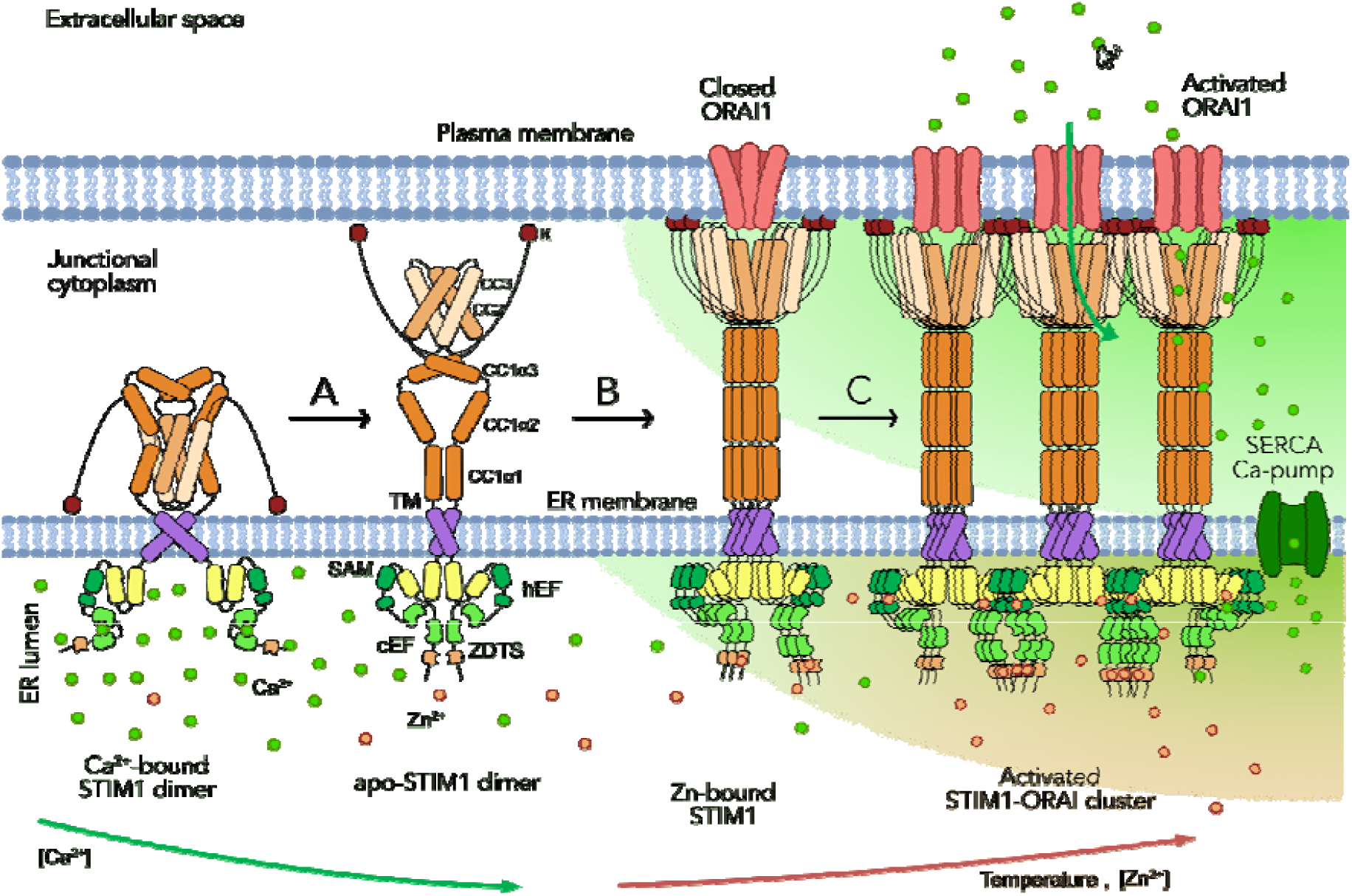
Schematic representation of temperature-dependent STIM1 activation by zinc ions. (A) A decrease in calcium concentration causes the dissociation of calcium ions from the luminal EF-hand domain of STIM1, triggering its partial unfolding. (B) The binding of zinc promotes STIM1 activation. (C) Zinc-linked STIM1 oligomers interact with the ORAI1 channel, facilitating calcium influx.

## Conclusion

Our findings reveal a previously unrecognized role of zinc in STIM1 activation and SOCE regulation. We demonstrate that the luminal domain of STIM1 harbors specific zinc-binding sites, with Cys49 and Cys56 playing a pivotal role in coordinating zinc ions. Structural and biophysical analyses indicate that zinc binding induces STIM1 dimerization and oligomerization, facilitating its clustering and interaction with ORAI1. Functional assays in HeLa and HEK cells confirm that disruption of the zinc-binding site impairs STIM1 clustering and calcium entry, underscoring the essential role of zinc in STIM1 activation. Furthermore, we provide evidence that intracellular zinc is required for ORAI1-mediated SOCE, in contrast to the inhibitory effects of extracellular zinc. These findings introduce an additional layer of complexity to SOCE regulation, highlighting the dynamic interplay between zinc and calcium in cellular signaling. Our study not only expands the current understanding of STIM1 activation but also suggests that zinc homeostasis could be a crucial factor in modulating calcium-dependent cellular processes.

## Materials and Methods

### Protein expression and purification

The luminal portion of human STIM1(amino acid 27 to 212) was cloned into pGEX-6P-1 without tag and overexpressed in Escherichia coli BL21 (DE3) strain. Bacterial cells were grown in LB with 100µg/mL ampicillin at 37°C, 150rpm. When the optical density at 600 nm reached 0.6, bacteria are induced thanks to 1mM isopropyl β-D-1-thiogalactopyranoside (IPTG) for 3h. Then the bacterial cells were pellet by centrifugation at 6500g for 15 min at 4°C and resuspended in a lysis buffer (50mM Tris HCl pH 7,5; 150mM NaCl). After a cycle of French press at 4 tones, lysis bacteria are incubated with lysozyme for 30 min at 4°C. On ice, the lysate is sonicated 5 times with cycles of 50s sonication, and 10s rest. After 20 min of centrifugation at 20,000g 4°C, pellet-containing STIM1 is re-solubilized at 12h in a 6M urea, 20mM Tris HCl pH 7.5, 1mM CaCl2 solution. The protein-containing urea suspension is dialyzed for 36h at 4°C against 2L of 20mM Tris Hcl pH 7.5, 0.1mM CaCl2. The dialysis buffer is changed every 12h. After 20 min of centrifugation at 20 000g, the supernatant containing STIM1 is loaded onto a 5 mL HiTrap DEAE Sepharose FF anion exchange column equilibrated in 20mM Tris HCl pH 7,5. STIM1 protein was eluted using 1M NaCl, 20mM Tris HCl pH 7,5 and dialyzed in 20mM Tris HCl pH 7.5, 0,1M NaCl. STIM1 protein concentration was measured spectrophotometrically at 280 nm using a molar extinction coefficient of 28 120 cm-1.M-1 and with bicinchoninic acid (BCA) dosage.

### Western Blot

Protein after purification was diluted 1:500 and loaded on wells of a 12% NuPAGE Bis-Tris for sodium dodecyl sulfate-polyacrylamide gel electrophoresis (SDS-PAGE) (Life Technologies), using Chameleon Duo as a size marker. Proteins are transferred onto a nitrocellulose membrane at 100V for 2h at 4°C. The membranes were blocked using Intercept Blocking Buffer (IBB-Li-Cor) for 30 min at room temperature (RT). Primary antibody (AntiSTIM1 N-terminal S6072 synthetic peptide corresponding to amino acids 61-74 of human STIM1, 1:500) was diluted in the IBB supplemented with 0.1% tween and incubated for 2h at RT. After three washes in PBS + 0.1% tween, the membrane was incubated with a secondary antibody (IR Dye 800CW donkey anti-rabbit, Li-Cor, 1:10,000) for 45 min at RT. The membrane was then washed thrice in PBS + 0.1% tween and once with PBS. Proteins were visualized using the LI-COR system.

### Mass spectrometry

Mass spectrometric measurements were performed on a Q Exactive instrument (Thermo Fisher Scientific, Bremen, Germany) equipped with the nanospray Flex source operating in infusion mode using the syringe pump at 1µl/min. The proteins were prepared at 60µM in 100mM MH4COOH with and without ZnCl2 or CaCl2 ions in excess.

### Isothermal titration calorimetry

The binding of ions to the EF-hand domain of STIM1 protein was studied on isothermal titration calorimeter iTC200 (MicroCal, United States) at 37°C in 50 mM Tris buffer, 1 mM TCEP at pH 7.5 as described previously ^35,36^ . The protein concentration in the calorimetric cell was 50 µM whereas the concentration of zinc and calcium ions in the titrating syringe was 0.5 mM. The volume of the injections was 0.5 µL before ions concentration reached an equimolar ratio with protein and then set to 2.5 µL. The dilution heat was measured by titration of the ion-containing buffer into the same buffer without the protein. The obtained curve was analyzed by MicroCal Origin 7.0 software using the models of the two types of binding sites. Therefore, the stoichiometry (N), constant (K_a_), enthalpy (ΔH), and entropy of binding (ΔS) were calculated from the standard thermodynamic equations.

### Differential scanning fluorimetry

The protein thermostability was measured in different concentrations of divalent ions in 50 mM Tris, at pH 7.5, 1 mM TCEP using a label-free fluorimetric analysis at Prometheus NT.Plex instrument (NanoTemper Technologies). NanoDSF grade capillaries were filled with the solution of EF-hand domain of STIM1 protein at concentration 37 µM (which corresponds to protein concentration in calorimetric call at the end of ITC titration) in the presence of either 1 mM Ca^2+^, or 1 mM Zn^2+^, or neither. The concentration of zinc or calcium ions varied from 18.5 to 370 µM. Capillaries were loaded onto the Prometheus NT.Plex instrument and heated from 20°C to 110°C with a 1 K/min heating rate. The excitation power was set at 100%. Unfolding transition points (T_m_) were determined from the first derivative of the changes in the emission wavelengths of the ratio of tryptophan fluorescence intensities at 350 nm and 330 nm (I_350_/I_330_), which were automatically identified by the Prometheus NT.Plex control software. With the same equipment, the aggregation temperatures (Tagg) were determined based on the first derivative of the temperature dependence of light scattering at 350 nm.

### Dynamic Light Scattering

Dynamic light scattering (DLS) experiments were carried out using a Zetasizer Nano ZS (Malvern Instruments) at 37°C. Replicate measurements were performed; each one consisting of 10-15 runs of 10 seconds. The scattering angle was 173°. The EF-hand domain of STIM1 was diluted to 50µM and centrifuged at 14000 rpm for 20 minutes. STIM1 size was determined in the absence or presence of increasing concentrations from 0.5 to 10mM of zinc or calcium ions. For the determination of the hydrodynamic diameter (Dh), the provided software uses the Stokes-Einstein relation to obtain the intensity averaged size distribution and the Mie theory to determine the volume size distribution which requires the viscosity and refractive index values of the dispersion medium and sample respectively. We used the dispersant viscosity of 0.6909 cP and protein refractive index of 1.450 (at 37°C) for both sizes.

### Surface Plasmon Resonance

SPR experiments were performed at 25°C with a Biacore T200 instrument (Cytiva) and using 10 mM HEPES-NaOH, pH7.4, 150 mM NaCl as running buffer. STIM1 was covalently immobilized on sensor chip CM5 (Cytiva) according manufacturer’s instructions. Briefly, two flow cells were activated by the injection of 35 µl of a 1:1 mixture of 50 mM N-hydroxysuccimide (NHS) and 200 mM N-ethyl-N’-(3-diethylaminopropyl)-carbodi-imide (EDC) at 5 µl/min. STIM1 was diluted at 2 µM in 10 mM acetate, pH 5.5, 20 µM CaCl_2_ and injected during the 420s at 5 µl/min over the experimental flow cell. CaCl_2_ was used to ensure that STIM1 remained monomeric during immobilization. The STIM1 immobilisation level was 87.8 fmoles/mm2. The remaining active sites were then blocked with 35 µl 1 M ethanolamine at 5 µl/min over the two flow cells. CaCl_2_, ZnCl_2_ and EDTA were serially diluted in the running buffer in the presence or not of STIM1 at 25 µM and injected over flow cells during 240 s at 10 µl/min. Sensorgrams with control flow cell (without immobilized STIM1) were systematically subtracted from those obtained with experimental flow cell (with immobilized STIM1). Sensorgrams obtained without STIM1 as analyte (only with Me^2+^or EDTA) were subtracted from sensorgrams obtained with STIM1 injection. Results are representative of 2 independent experiments.

### Transmission Electron Microscopy

Protein solution at 10µM, with or without 160µM CaCl2 or 80µM ZnCl2, was incubated on 200-mesh copper grids for 3 min. The grids were previously heated to 37°C and all the preparation was done at 37°C. After removing the excess of solution with filter paper, 2% uranyl acetate was applied for 45s and then blotted off. The grids were observed with G2 FEI Technai 200KV transmission electron microscope. Images were made with a 2K Veleta camera.

### Cell culture, transfection, and fluorescence microscopy

HeLa and HEK293 cells were cultured in Dulbecco’s Modified Eagle Medium (DMEM) with high glucose, GlutaMAX Supplement, and pyruvate (Gibco, #31966-021), supplemented with 10% Fetal Bovin Serum (Gibco, #5209402) and 1% Penicillin Streptomycin (Gibco, #15070-063). Cells were maintained at 37°C in a humidified atmosphere containing 5% CO□. For fluorescence microscopy, cells (50,000 cells per well) were seeded onto sterile 25mm coverslips placed in 6-well plates. Transfection with Lipofectamine 3000 (Invitrogen, #L3000015) was conducted 48 hours post-seeding using 1 or 2 µg of full-length STIM1-EGFP WT or Cys-STIM1 (mutated STIM1 C49A, C56A, H57A) in the pEGFP-N1vector. The transfection medium was replaced after a few hours, and the cells were cultured for an additional 48 hours before experimentation.

Cells were observed using an Axio Observer.Z1/7 microscope (ZEISS) under 5% CO□ and 37°C conditions. Imaging was performed using a 63X/1.20 W objective for STIM1 activation or a Fluar 20X/0.75 M27 objective for the Fura-2 experiment and a Hamamatsu Flash 4 camera.

### STIM1 activation and Fura-2 Fluorescence Experiments

Transfected HeLa cells with STIM1-EGFP WT or Cys-STIM1 were observed in HBSS medium (138 mM NaCl, 5.3 mM KCl, 0.34 mM Na_2_HPO4; 0.44 mM KH_2_PO_4_; 4.17 mM NaHCO_3_; 4mM Mg²□, 1.3 mM CaCl_2_ at pH 7.4) under 5% CO□ at 37°C. Images were made before and after the addition of 1µM thapsigargin (TG) to induce calcium store depletion and STIM1 activation.

Transfected HEK293 cells were loaded with 5 µM Fura-2-red, an AM dye (Invitrogen, #F3021) dissolved in DMSO (Invitrogen, #D12345) and prepared in HBSS medium supplemented with 1% Pluronic F-127 (Invitrogen, #P3000MP). Cells were incubated in the dark at 37°C for 45 minutes. Then, they were washed with calcium-free HBSS (composition as above, but without CaCl□) and coverslips were mounted in the microscope chamber in calcium-free HBSS.

Dual-excitation wavelengths were employed with filter sets: Cy5 illumination at 401–445 nm for 430 nm excitation, and Cy5 at 450–488 nm for 475 nm excitation, with emission detection at 680 nm (659–701 nm filter). To induce calcium store depletion, 0.1 mM TG (Sigma, #T9033) was added, and calcium flux measurements were captured for 6 minutes at 5-second intervals. Store-operated calcium entry (SOCE) was assessed by adding 2 mM CaCl□ to the TG-containing medium, with measurements every 10 seconds for 3 minutes. To determine maximal calcium entry capacity, 1 µM ionomycin was applied to permeabilize the cells, followed by 10 mM CaCl_2_. Cytosolic calcium concentration (□Ca²□] □) was measured by selecting ROIs from the recorded images.

### ML protein co-folding

The structure of the EF-STIM1 dimer complex with zinc was modeled using the ML approach of Chai-1 protein co-folding^37^. Chelator quadruples were combinatorially constructed for residues 23, 30, and 31, and distance constraints characteristic of zinc chelation by the corresponding amino acid quadruple were created for all 15 combinations. Two EF-STIM1 molecules and three zinc ions were co-folded with these constraints. The selection of successful structures was based on the formation of zinc chelation by the selected residues.

## Declarations

### Authors’ contributions

Conceptualization, M.B. and P.O.T.; methodology, M.B. and P.O.T.; investigation, V.E.B., B.A., S.B.,C.V., and S.C.; writing—original draft, M.B., B.A., and P.O.T.; writing—review & editing, M.B., B.A., and P.O.T.; funding acquisition, M.B.; resources, F.D.; supervision, M.B. and P.O.T.

### Competing interests

Authors declare no competing interests.

## Funding

This work was supported by an AFM-Telethon grant under the strategic program Mothard.

## Acknowledgments

Microcalorimetry and nanoDSF experiments were performed at PINT (Plateforme Interactions moléculaires Timone), Marseille, France. Prometheus NT.Plex instrument was acquired with the financial support of Région PACA (APR-EX 2016 Neuro-Plasma). We thank the PiCSL-FBI core facility (IBDM, AMU-Marseille), members of the France BioImaging national research infrastructure, in particular Nicolas Brouilly, who performed the electron microscopy experiments. We also thank Dr. Olivier Dellis for his insightful comments and reflections, and Dr. Alexandra Salvi for the design of the expression plasmids.

## Resource availability

All data available upon request.

## Abbreviations

SOCE: store-operated calcium entry
ORAI: calcium release-activated calcium channel protein
CRAC: calcium release-activated channels
STIM: stromal interaction molecule
ITC: isothermal titration calorimetry
DLS: dynamic light scattering
DSF: differential scanning fluorimetry
EF-STIM1: 27-212 a.a. fragment of STIM1 luminal domain

## Notes

### Competing Interest Statement

The authors have declared no competing interest.

